# Two-Step Mechanism of Bruton’s Tyrosine Kinase Membrane Recruitment and Activation

**DOI:** 10.1101/2025.10.28.685231

**Authors:** Rachel A. McAllister, Amy L. Stiegler, Keerthana Chari, Meera Chari, Moitrayee Bhattacharyya, Kallol Gupta

## Abstract

Peripheral membrane proteins (PMPs) are critical mediators of signaling cascades initiated via activation of cell-surface receptors. Their functions rely on their innate ability to interact with membranes dynamically in response to rapidly changing cellular conditions. This membrane recruitment may occur via high-affinity interactions with specific lipids or transient, low-affinity membrane interactions. These weak and dynamic interactions that are critical regulators of protein function are challenging to capture. Taking Bruton’s Tyrosine Kinase (BTK), a non-receptor tyrosine kinase essential for B cell activation, as an example, we demonstrate a native mass spectrometry (nMS) platform to understand the recruitment of PMPs by directly studying it from lipid bilayers customized to target membranes. Our data demonstrates that BTK recognizes phosphatidylserine (PS) through sites distinct from phosphatidylinositol (3,4,5) phosphate (PIP_3_) binding. We show PS-bound BTK retains PIP_3_ binding simultaneously via high-affinity sites, exhibiting PIP_3_-independent basal membrane recruitment of BTK. Biochemical assays show that this PS-mediated recruitment sensitizes BTK to PIP_3_-mediated activation under near-physiological concentrations of PIP_3_. Thus, we propose a two-step model for BTK membrane recruitment and activation. A low-affinity interaction with high-copy number PS enables plasma membrane recruitment of BTK and increases its membrane-bound concentration. Upon B-cell activation, this pre-recruited, membrane-bound BTK population localizes to PIP_3_-rich domains through electrostatic gliding along the membrane driven by low-affinity PS and high-affinity PIP_3_ binding. This indicates a cooperative mechanism where PS can amplify B-cell signaling through increased membrane-bound BTK concentration. Our work demonstrates a general model of PH-domain-containing protein regulation by weak protein-lipid interactions, which can be extended to many other PMPs.

## Introduction

Signaling pathways within the cell that initiate from organellar membrane receptors often rely on downstream peripheral membrane proteins to propagate the signals into cellular actions. This includes the MAPK pathway that regulates cell growth, B-cell receptor pathway that regulates immune response, and mitochondrial fission-fusion dynamics that regulate mitochondrial health (1–3). A significant subset of peripheral membrane proteins conditionally associates with membranes via specific recognition of signaling lipids such as phosphatidylinositol (3,4,5) phosphate (PIP_3_) (4–7). Among the most well-studied peripheral membrane protein domains is the Pleckstrin Homology (PH) domain, the 11^th^ most common domain in the human proteome (8). A key PH-domain containing peripheral membrane protein that plays an essential role in B Cell Receptor (BCR) signaling is Bruton’s tyrosine kinase (BTK) (9, 10). BTK is a Tec family kinase whose PH domain has an adjoining unique zinc finger motif required for structural stability (Fig. 1A) (11, 12). Upon antigen stimulation, the BCR undergoes clustering, leading to SYK/LYN-mediated phosphorylation of the immunoreceptor tyrosine-based activation motifs (ITAMs) on the associated immunoglobulins CD79a and CD79b (13, 14). This process initiates the downstream activation of Phosphatidylinositol (3) Kinase (PI3K), which phosphorylates membrane lipid phosphatidylinositol (4,5)-bisphosphate (PIP_2_) to form phosphatidylinositol (3,4,5)-trisphosphate (PIP_3_) at the plasma membrane (15, 16). Previous studies have shown that PIP_3_ binding leads to activation of BTK, leading to auto-phosphorylation on Tyr223 (12, 17). In the cytoplasm, BTK exists as an autoinhibited monomer with the SH2 and SH3 domains compact against the distal face of the kinase (Fig. 1A) (18, 19). On the membrane surface, PIP_3_ binding is suggested to directly compete with one autoinhibitory conformation of the PHTH domain in the full-length kinase and stabilize dimeric interfaces (20). This ability to relieve autoinhibition and stabilize BTK interactions is critical for BTK *trans-*autophosphorylation within the cell (18, 21). Additionally, SYK or Src-family kinases, parallelly activated downstream of the BCR, phosphorylate Tyr551 in the kinase domain activation loop of BTK, further increasing its catalytic activity (13, 22). Activated BTK phosphorylates and activates phospholipase-Cγ2 (PLCγ2), which cleaves PIP_2_ to produce diacylglycerol (DAG) and soluble secondary messenger inositol triphosphate (IP_3_) (23, 24). IP_3_ binds to the IP_3_-receptor in the endoplasmic reticulum (ER), releasing calcium and activating Nuclear Factor of Activated T cell (NFAT) and Nuclear Factor-kappa B (NF-κB) pathways, leading to internalization of the antigen and activation of the B Cell (25–27). Due to the central role of BTK in adaptive immunity, mutations within its canonical PIP_3_ binding pocket have been shown to cause X-linked agammaglobulinemia (XLA), a severe immunodeficiency syndrome (Fig. 1B) (28, 29). A key step in the BCR activation is the plasma membrane recruitment and activation of BTK (12, 21, 30, 31). While other modes of activation in solution, such as activation via the inositol hexakisphosphate (IP_6_) through a novel binding site have been recently uncovered (18), the current mechanistic understanding is that PIP_3_ alone is responsible for both the recruitment of BTK to the plasma membrane and its activation in the membrane. The central hypothesis of this study is that BTK can simultaneously interact with other abundant plasma membrane inner leaflet lipids, in addition to PIP_3_, using binding sites distinct from those used for PIP_3_ binding. We propose that these likely weaker interactions promote PIP_3_-independent recruitment of BTK to the plasma membrane in an inactive state, increasing its local concentration and enabling BTK to scan along the quasi-two-dimensional inner leaflet of the plasma membrane for PIP_3_. Upon generation of PIP_3_, this membrane pre-localized pool enables a stronger activation response, leading to a synergistic mechanism by which orthogonal weak affinity lipid interactions can amplify PIP_3_-mediated BTK activation and subsequent signaling. Indeed, while PH domains were initially thought to primarily to recognize PIPs, in recent years studies have shown a broad range of functions, including work on the isolated PH domains of GRP1, ASAP1, and PDK1 showing background electrostatic lipids induced higher membrane residence (32–35). These studies all suggest that a broader mechanism of coincidence detection of negatively charged phospholipids on the target membrane may play a significant role in recruiting peripheral membrane proteins for activation. Establishing this dual-mode lipid interaction demands experimental avenues that offer a dynamic detection range and molecular resolution to simultaneously detect both tightly and weakly bound lipids and unambiguously determine their identities and stoichiometry. Furthermore, it demands technical abilities to perform these analyses on formed target protein lipid complexes directly from a membrane environment that can be customized to a target physiological membrane. We have recently developed a native mass spectrometry (nMS) approach that enables the study of integral membrane proteins from customizable proteoliposomes, allowing for a quantitative understanding of their hierarchical organization and lipid binding directly from lipid bilayers mimicking different organellar membrane contexts (36, 37). In the current work, we further optimize and extend this platform to study how specific lipids regulate the recruitment of peripheral membrane proteins to organellar membranes, taking BTK as the target protein. A key advantage of our nMS is its unmatched molecular resolution to detect and identify lipids that are specifically bound to target proteins—both individually or together—along with their binding stoichiometries, directly from complex lipid bilayers (36, 37). The general sensitivity of nMS enables further detection of even a minor fraction of protein bound to a lipid, allowing for application to a wide and dynamic range of protein lipid interactions (38–42). Applying this lipid vesicle nMS platform, we discovered that BTK can get recruited to the membrane surface through a weak affinity interaction with phosphatidylserine (PS), independent of PIP_3_. We further established that this low-affinity interaction with PS, a highly abundant lipid in the plasma membrane inner leaflet, takes place via sites independent of the canonical PIP_3_ binding site or the recently proposed peripheral site. Using *in vitro* kinase assays, we further show that this PIP_3_ independent PS mediated recruitment leads to an increase in the degree of BTK autophosphorylation in the presence of physiologically relevant amount of PIP_3_. This builds a two-step recruitment and activation mechanism of BTK, where PIP_3_ and PS work in a synergistic manner. Beyond BTK, the work also presents a methodological platform to understand membrane recruitment of peripheral membrane proteins with precise molecular detail.

**Figure 1.**
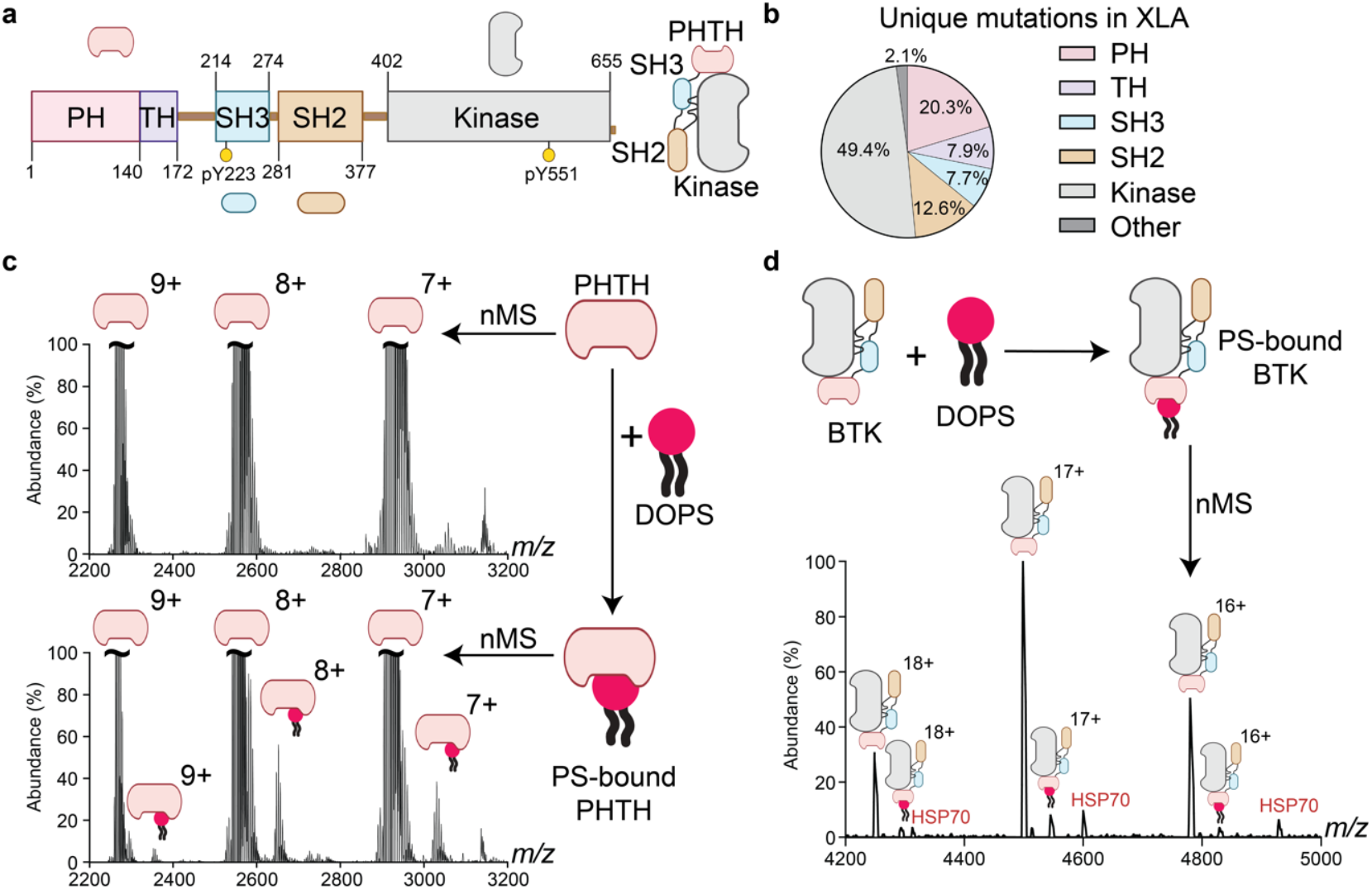
Structural features, XLA mutations, and PS binding of BTK: **(a)** Domain architecture of BTK showing the N-terminal PH and Tec Homology (TH) domain which contains a zinc finger motif, followed by an Src module composed of: an Src-homology 3 (SH3) domain, Src-homology 2 (SH2) domain, and a C-terminal kinase domain. Yellow circles denote major tyrosine phosphorylation sites Tyr223 and Tyr551. The structural model shows the autoinhibited state, in which the PHTH domain primarily folds against the kinase domain but can adopt a range of conformations (18, 20, 43). **(b)** Percentage distribution of unique X-linked agammaglobulinemia (XLA)-associated missense mutations mapped to each domain within BTK via *BTKbase* web-repository (29). **(c)** Native mass spectra of the isolated PHTH domain **(top)** and in the presence of 18:1-18:1 phosphatidylserine (DOPS) **(bottom)**. Charge states are labeled and PS-bound peaks are marked. Observance of PS-bound peaks to all charge state peaks confirms the binding of PS to BTK. Spectra are normalized to 10% of major species in each spectra and repeated in triplicate. **(d)** Native mass spectra of the full-length BTK in the presence of DOPS. Minor purification impurities due to the presence of Heat Shock Protein 70 (HSP70) are annotated with a red HSP70. Like the isolated PHTH domain, the full-length BTK also binds to PS. The spectra were repeated in triplicate.

## Results

Our first goal was to determine if BTK can bind to other plasma membrane lipids beyond PIP_3_. To address this, at first, we utilized just the membrane binding PHTH domain of BTK and studied independent lipid binding in solution using nMS. By directly incubating the PHTH domain with different lipids independently and subjecting it to nMS, we found that the PHTH domain is capable of binding PS (Fig. 1C), a negatively charged phospholipid that is highly abundant on the plasma membrane. PS is primarily localized to the inner leaflet, accounting for ~1/5^th^ of inner leaflet lipids (44–48). In contrast, no binding was detected for phosphatidylethanolamine (PE), another prominent plasma membrane lipid enriched on the inner leaflet, indicating the specificity of the protein for the anionic serine head group over the more zwitterionic PE head group (Fig. S1A-D) (44, 48). In the full-length autoinhibited conformation of the protein, the PH domain is sandwiched against the N-lobe of the kinase, yet explores a range of conformations, leading to minor heterogeneity in the auto-inhibited state (Fig. 1A) (18–20). Due to this potential occlusion of PHTH domain surfaces, we wanted to confirm that full-length BTK can also recognize PS. As shown in Figure 1D, full-length BTK also binds to PS, indicating that binding to PS is not abrogated by the PHTH domain conformation in the full-length protein. Finally, we aimed to demonstrate that this PS-BTK interaction can also occur in the presence of a bilayer, within the lipid-rich milieu that comprises the collective membrane environment. This is a critical consideration when studying protein-lipid interactions. While a target protein may bind a specific lipid in isolation, such interactions may not persist in the context of a complex physiologically relevant lipid bilayer(49). In these environments, other abundant lipids could potentially compete for binding, disrupt, or modulate the interaction (36, 37). Therefore, assessing lipid binding within native or native-like lipid mixtures is essential for accurately capturing biologically meaningful interactions.

To address this, we incorporated our recently published nMS method that enables the study of integral membrane protein-lipid interactions directly from customizable liposomes (36, 37). Here we adapted this approach to peripheral membrane proteins, where gentle ionization and in-source activation disrupts the vesicles enough to resolve direct protein-lipid complexes (Fig. 2A). First, from liposomes containing phosphatidylcholine (PC), PE and a minor amount PS – we observe a robust and exclusive PS binding peak (Fig. 2B). Next, we customized the lipid composition of the liposome to more closely resemble the lipid composition of the inner leaflet of the plasma membrane, the leaflet encountered by cytosolic peripheral membrane proteins. Typically, the inner leaflet is expected to be devoid of cholesterol and sphingomyelin, which primarily reside on the outer leaflet, and high PS (47, 48). As shown in Figure 2C, even in the plasma membrane inner leaflet-like liposome containing 7 different phospholipid species, PS binding is observed as the primary adduct. The ability to detect robust PS binding to BTK incubated with these custom lipid bilayers indicates the ability of PS in an inner leaflet-like bilayer to recruit BTK to the membrane surface.

**Figure 2.**
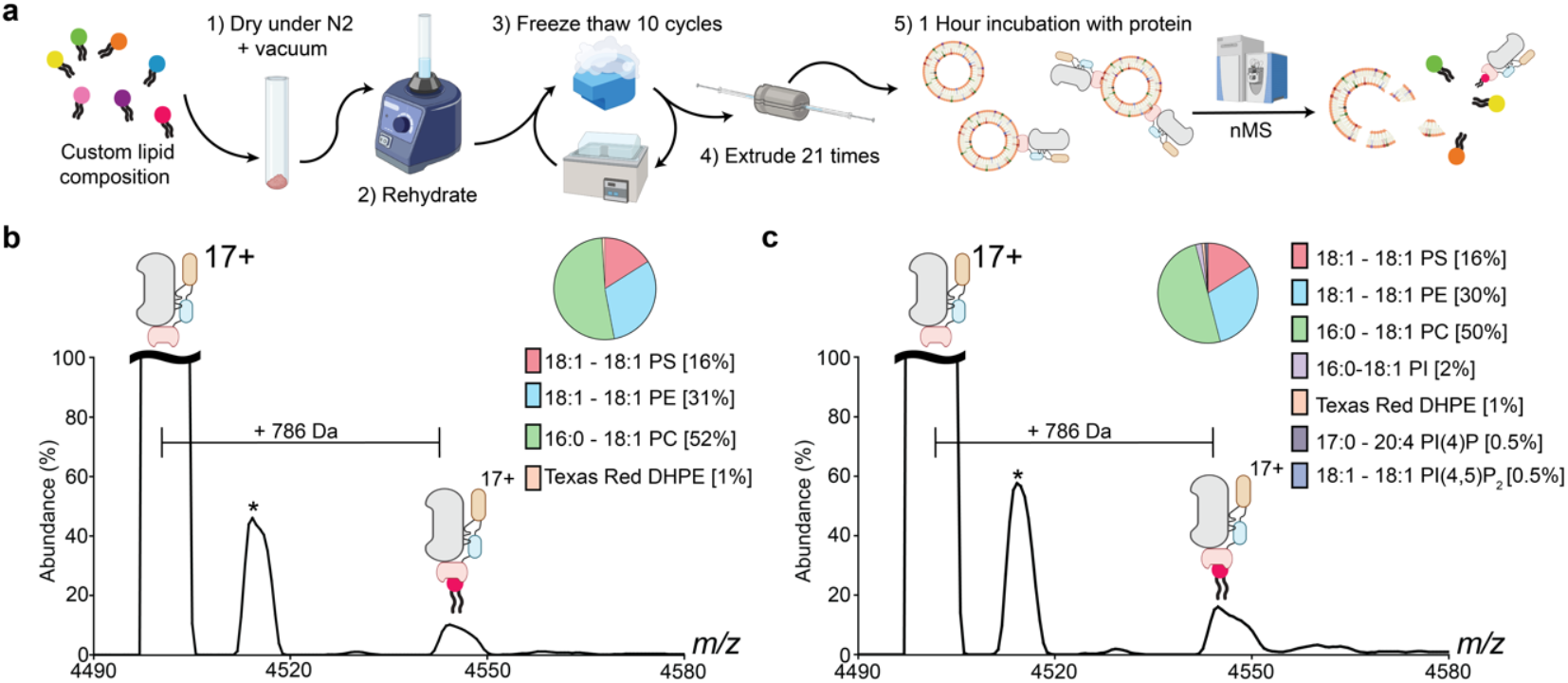
Lipid binding to BTK directly from the plasma membrane inner leaflet-like liposome. **(a)** Flow chart of peripheral membrane protein proteo-liposome preparation. Lipids in chloroform are dried under a nitrogen flow for 30 minutes, following a minimum of two hours under vacuum, to remove residual organic solvent. Aqueous nMS buffer is added to the dried lipid film for vortex rehydration prior to ten cycles of alternating liquid nitrogen and a 50 °C water bath, followed by extrusion with 100 nm pore membranes using an Avestin Liposofast basic. The homogenous vesicles are then incubated with protein prior to nMS, where the protein can be ejected from these vesicles with bound lipid still attached. **(b)** Mass Spectra of full-length BTK in the presence of PC/PE/PS vesicles. Lipid vesicle composition is shown on the right. As annotated, direct and exclusive binding of PS to BTK could be observed from the liposome. Spectra were normalized to 10% of the apo peak and repeated in triplicate. **(c)** Mass Spectra of full-length BTK in the presence of inner-leaflet plasma membrane-like vesicles. Lipid vesicle composition shown on the right. Exclusive binding to PS to BTK was still observed, despite the presence of other inner-leaflet anionic lipids at physiological ratios. Spectra were normalized to 10% of the apo peak and repeated in triplicate. In both spectra, the * denotes a minor impurity of 241 Da addition that corresponds to a covalent phosphogluconylation of BTK. A noted and established artifact of overexpression of membrane-binding domains/proteins in E. coli (7).

To corroborate the nMS-based observations, we performed fluorescence polarization (FP) assays in solution. Fluorescent nitrobenzodiazole (NBD) – labeled PS, modified on the fatty acyl chain to preserve headgroup sterics, was kept at a fixed concentration while the PHTH domain concentration was varied. Excitation at 485 nm of polarized light and emission detection at 528 nm were used to monitor NBD fluorescence. In the absence of binding, the small, fast tumbling PS exhibits low polarization as the emitted light is largely depolarized due to rapid Brownian motions during the NBD excitation lifetime (50). Upon addition and binding to the bulkier PHTH domain the rotational speed is decreased and less light is depolarized. As shown (SI. Fig. 2A), this increases the polarization read-out, providing direct solution-phase evidence of BTK-PS binding. We further observe a PHTH-concentration-dependent increase in polarization, consistent with binding of the PHTH domain to PS (SI. Fig. 2B).

Next, we asked whether the PS binding to BTK can occur independently of PIP_3_ binding, as both PS and PIP_3_ are negatively charged lipids that may recognize similar positively charged residues on proteins. To test this, we employed dual charge reversal mutations at the canonical PIP_3_ binding site formed by Lys 12, Arg 28, Asp 24 and Tyr 39 and performed nMS of this R28C/N24D mutant BTK, which has been shown to abolish the PIP_3_ binding to BTK (18, 28, 30). nMS-based lipid binding studies on this mutant reveal that the mutant is still capable of binding to PS (SI. Fig. 3A-C). We next tested a recently discovered secondary binding site for Inositol Hexakisphosphate (IP_6_), coordinated through Arg 52, Lys 36, Lys 49 and Tyr 40 (18). nMS of K49S/R52S mutant PHTH also shows no significant drop in the PS binding ability over the wild type protein (SI. Fig. 4A-C). Together, these data indicate that BTK can bind to PS independent of the specific PIP_3_ binding sites.

To further establish the independent binding mechanism, we performed a series of dual binding experiments involving full-length BTK, PIP_3_, and PS. First, to unambiguously establish that such dual binding can occur directly from a membrane environment, we performed nMS of BTK in the presence of PS-containing liposomes, where we added IP_4_, the functionally active headgroup of PIP_3_ (SI. Fig. 5). Prior work has established that IP_4_ adopts the same binding pose as PIP_3_ at the canonical site (51). Additionally, at a steady concentration of BTK and PS, we increased IP_4_ concentration, and the resultant solution was subjected to nMS analysis. The apo and lipid/IP_4_ BTK peaks were quantified via the area under the curve for the predominant charge state. As shown in Figure 3A, and quantified in Figure 3B-C, the addition of IP_4_ leads to a clear observation of IP_4_-bound BTK and critically, no significant loss in PS binding was observed. More importantly, as the concentration of IP_4_ in solution increases, a distinct peak at m/z 4574 arises, corresponding to the mass of BTK bound simultaneously to IP_4_ and PS. This indicates that PS-bound BTK can simultaneously engage and bind the PIP_3_ head-group. Finally, we confirmed the dual lipid-binding capability of BTK by incubating PHTH domain with both PS and PI(3,4,5)P_3_. nMS of this mixture identified a characteristic peak that corresponds to the simultaneous PIP_3_+PS bound state (Fig. 3D). This peak was absent in controls with BTK incubated with only PS or PIP_3_ (SI Fig. 6). Together, these data unambiguously establish that BTK can simultaneously bind PS while engaging with PIP_3_ through the canonical binding site. Interestingly, the plasma membrane is composed of approximately 10% PS, with the majority localized to the inner leaflet, constituting about 15-20 mol% of inner leaflet lipids (44, 46). For cytosolic peripheral membrane proteins like BTK, which exclusively interact with the inner leaflet, this lower-affinity interaction with an abundant lipid like PS can facilitate dynamic membrane exploration. Specifically, this PS-dependent interaction can position BTK favorably on the membrane surface, allowing its PH domain to efficiently search for and engage high-affinity activating ligands, such as PIP_3_. Importantly, this PS-mediated basal membrane association should also inherently elevate the local BTK concentration at the plasma membrane independently of PIP_3_ availability.

**Figure 3.**
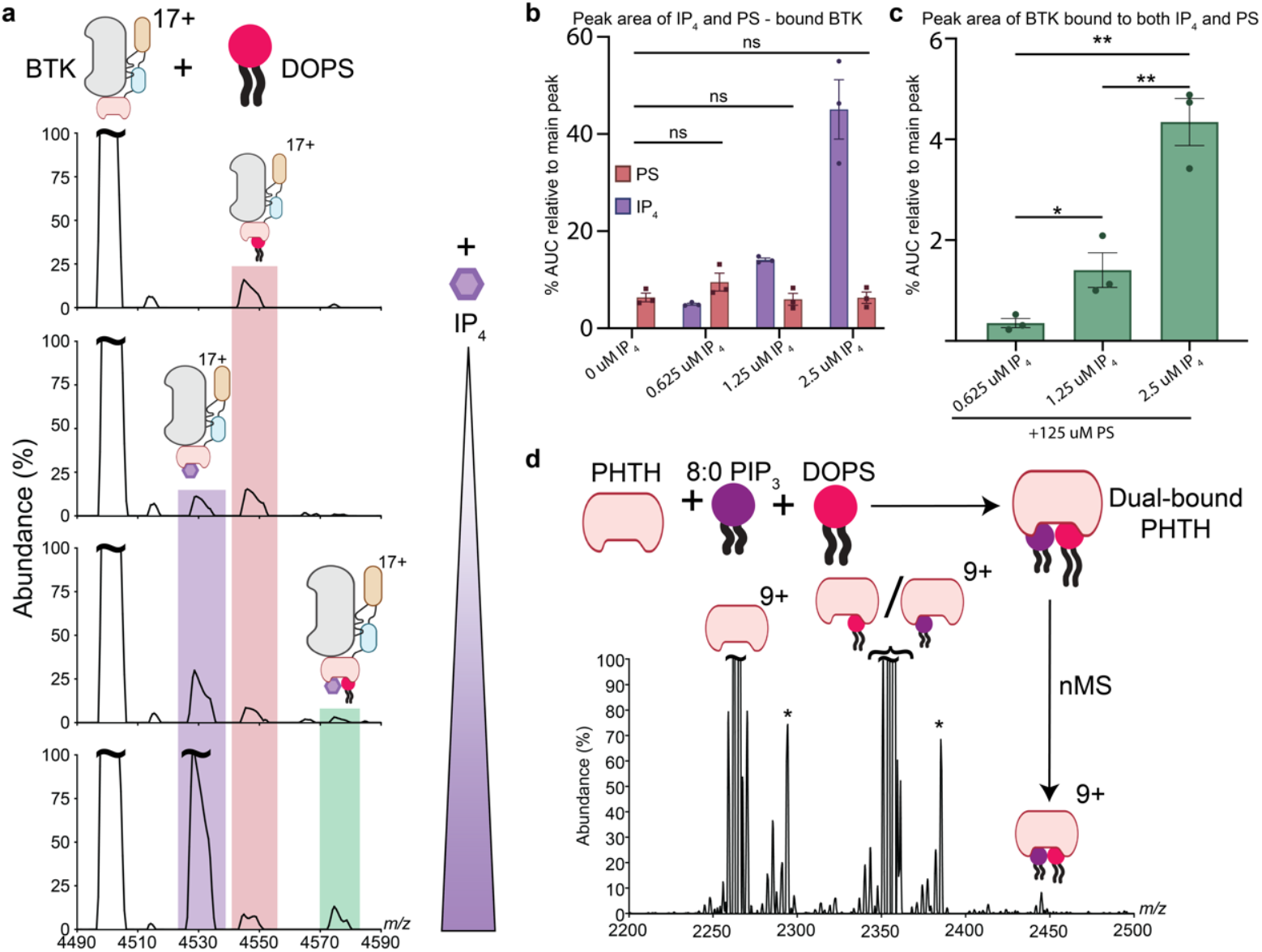
Ability of BTK to bind both PS and PIP_3_/IP_4_ simultaneously. **(a)** Expansion of charge state 17 of nMS of full-length BTK after incubation with DOPS. Top to bottom: Increasing amounts of inositol tetrakisphosphate (IP_4_) are added. BTK is kept at 2.5 μM, PS at 125 μM, and IP_4_ concentration (top to bottom: 0 μM, 0.625 μM, 1.25 μM, and 2.5 μM). Peak abundance is normalized to 50% of the charge state 17 peak. As highlighted in a light fuchsia, all conditions produce a PS-bound peak. Incubation with IP_4_ yields an IP_4_-bound peak highlighted in purple. Additionally, increasing IP_4_ concentration yields a peak, highlighted in green, that corresponds to the BTK+PS+IP_4_ species. **(b)** Plot of the peak area of the PS and IP_4_-bound peaks obtained from (a), highlighted in red and purple, respectively. The area under the curve (AUC) was calculated for each as a percentage of the apo charge state peak (mean ± sem, n = 3). Statistics calculated using an unpaired, nonparametric t-test on Graphpad Prism. Statistical significance defined as p>0.05, ns; p< 0.05 as *, p<0.01 as ** and p<0.001 as ***. **(c)** Plot of the peak area of the peak at m/z = 4574 corresponding to dual IP_4_ and PS-bound BTK (highlighted in light green) (mean ± sem, n =3). Statistics calculated using an unpaired, nonparametric t-test on Graphpad Prism. **(d)** Spectra of the wild-type PHTH domain upon incubation with both 8:0 PIP_3_ and PS reveal a unique peak at m/z 2445, which corresponds to the dual-bound species.

Given that BTK activation critically involves trans-autophosphorylation, this PS-mediated increase in local BTK concentration on the plasma membrane is likely to facilitate autophosphorylation and enhance BTK activation, particularly under conditions of limited PIP_3_ availability. To assess the influence of PS-dependent membrane recruitment on BTK activation, we utilized an *in vitro* BTK autophosphorylation assay (18). We set up BTK phosphorylation assays in the presence of liposomes containing various concentrations of PS and PIP_3._ Briefly, BTK was incubated with custom liposomes before the introduction of ATP and MgCl^2+^ to initiate reaction. The reaction was stopped after 5 minutes by adding SDS buffer + EGTA and boiling the sample (Fig. 4A). The extent of BTK autophosphorylation was measured and quantified by western blots using BTK phosphoTyr223 antibodies and normalized with respect to total BTK concentration. Previous work has established that PIP_3_ is necessary and sufficient for activation of trans-autophosphorylation of BTK on the membrane (28, 52). Aligning with this observation, PS and other phosphatidylinositol-containing liposomes (PIP or PIP_2_) failed to activate BTK by themselves in the absence of PIP_3_ (SI Fig. 7), affirming that PIP_3_ is necessary for BTK activation. This aligns well with previous mechanistic work that demonstrates how stable membrane binding of PIP_3_ facilitates BTK autophosphorylation (12, 30). Next, BTK autophosphorylation assays conducted with liposomes containing high levels (5%) of PIP_3_ showed autophosphorylation, although no additional influence from increasing PS concentrations in the liposomes could be observed (Fig. 4b). However, under 10-fold lower PIP_3_ concentration (1-0.5%), incremental increases in PS concentration led to a near-linear enhancement of BTK autophosphorylation (Fig. 4C-D). Notably, the magnitude of this PS-mediated amplification of autophosphorylation inversely correlated with PIP_3_ abundance – reduction in PIP_3_ concentration heightened the impact of PS in the liposomes. Strikingly, at 0.5% PIP_3,_ we note 2.77-fold increase in kinase autophosphorylation going from 0% PS to 20% PS (Fig. 4D). Based on our experimental data, we constructed a simple mathematical model to understand how PS-mediated membrane recruitment can increase the rate of BTK autophosphorylation (See Methods for details). For this, we leveraged a previously established framework and further modified it to approximate how the effective concentration of BTK on the liposome surface would change as a result of PS at low PIP_3_ concentrations (18). This simple and approximate model predicts approximately 1.97-fold increase in the autophosphorylation rate at 0.5% PIP_3_ going from 0 to 20% PS (SI Fig. 8). This closely agrees with experimentally derived change in BTK autophosphorylation, further establishing that increase in *trans*-autophosphorylation is a direct representation of an increase in effective concentration in the presence of the liposomes. Together, these results are in agreement with our nMS-based observations and a two-step model that low-affinity interactions with abundant PS enhance local BTK concentrations at the membrane, consequently promoting the ease and extent of BTK autophosphorylation under lower concentrations of PIP_3_.

**Figure 4.**
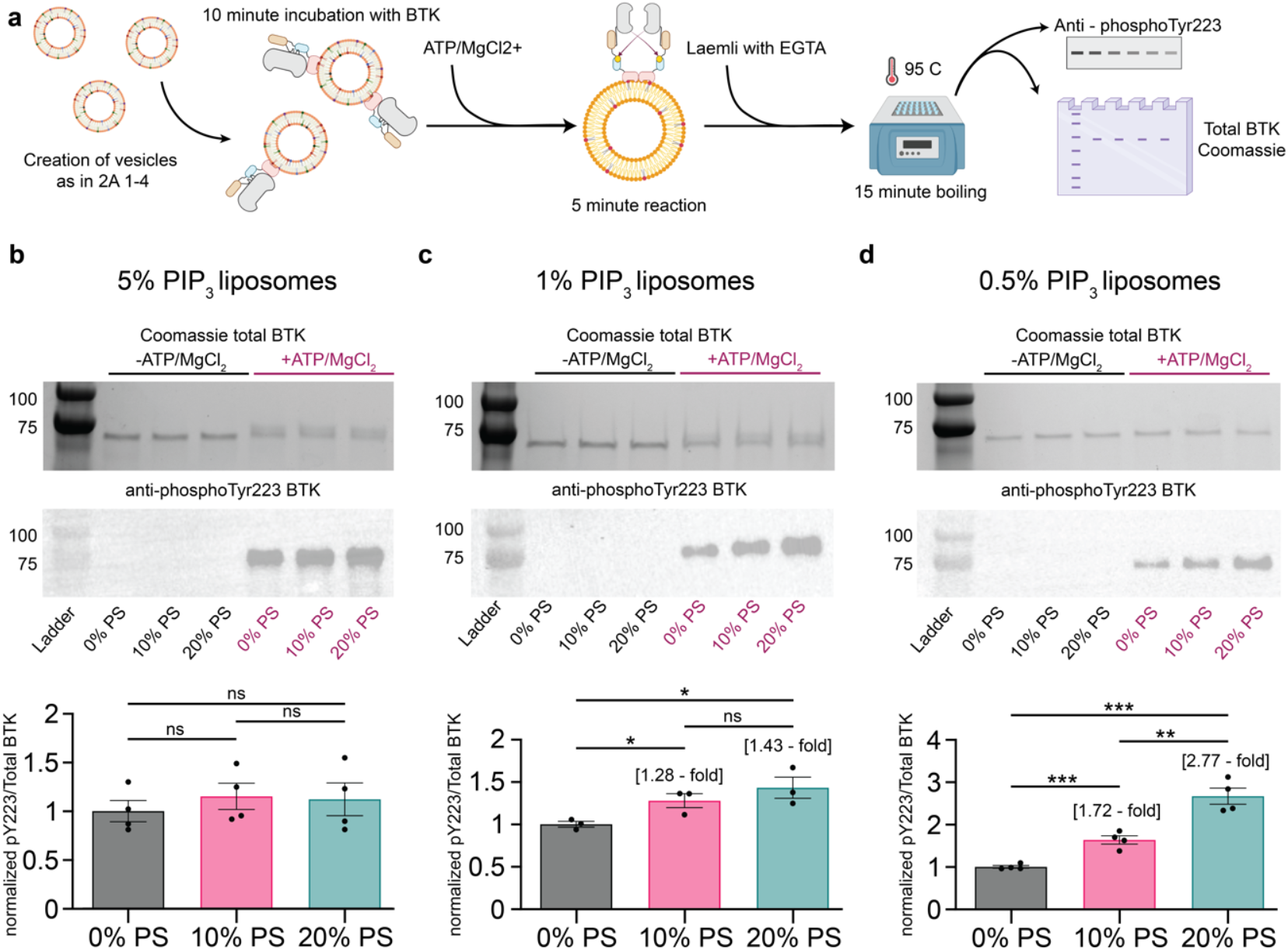
Influence of PS on the trans-*auto*phosphorylation of BTK. **(a)** Schematic depicting the kinase assay. Briefly, liposomes are created as in Fig 2A. After a 10-minute pre-incubation with BTK, kinase reaction buffer is added and reaction proceeds for 5 minutes. The reaction was stopped by the addition of Laemli w/ EGTA and boiling. For more details, see methods. All conditions normalized to the mean of 0% PS for their respective PIP_3_ concentrations and representative westerns generated as a merge between colorimetric (ladder) and chemiluminescence. **(b) (top)** Representative Coomassie stain of 5% PIP_3_ liposomes to demonstrate total Btk concentration in both the negative control in the absence of ATP and MgCl_2_ as well as in the reaction conditions. **(middle)** Representative anti-phosphoTyr223 BTK western. **(bottom)** Plot of the quantified and normalized phosphoTyr223/Total BTK ratio (mean ± sem, n =4). No change in extent of phosphorylation observed despite changing PS concentrations. Statistical significance defined as p>0.05, ns; p< 0.05 as *, p<0.01 as ** and p<0.001 as ***. **(c) (top)** Representative Coomassie stain of 1% PIP_3_ liposomes to demonstrate total Btk concentration in both the negative control in the absence of ATP and MgCl_2_ as well as in the reaction conditions. **(middle)** Representative anti-phosphoTyr223 BTK western. **(bottom)** Plot of the quantified and normalized phosphoTyr223/Total BTK ratio (mean ± sem, n =3). Increases in relative phosphorylation from 0%PS to 10% PS or 20% demonstrate an increase in BTK activation in the background presence of anionic PS. **(d) (top)** Representative Coomassie stain of 0.5% PIP_3_ liposomes to demonstrate total Btk concentration in both the negative control in the absence of ATP and MgCl_2_ as well as in the reaction conditions. **(middle)** Representative anti-phosphoTyr223 BTK western. **(bottom)** Plot of the quantified phosphoTyr223/Total BTK ratio (mean ± sem, n =4). A 0.5% PIP_3_ concentration, a stark increase in autophosphorylation level is observed upon increasing the PS concentration, demonstrating a near linear increase in BTK activity with regards to increase PS concentration.

## Discussion

Our findings reveal a novel interaction between BTK and PS, an abundant negatively charged lipid predominantly localized on the inner leaflet of the plasma membrane. Importantly, this interaction occurs at sites independent from the canonical PIP_3_-binding pocket of BTK. This allows BTK to simultaneously engage with PS and dynamically explore the membrane surface for high-affinity interactions with PIP_3_. We confirmed this through mutational analyses that selectively disrupted PIP_3_ binding without impairing PS interaction. Given the very low physiological abundance of PIP_3_ (<1 mol %) and its short life span, the observed low-affinity and high-copy number PS interactions can substantially elevate local BTK concentration and facilitate rapid, dynamic membrane scanning by the PHTH domain for PIP_3_. The biological significance of this electrostatic interaction is highlighted by our kinase assays, where it enhances membrane-localized trans-autophosphorylation, thereby lowering the threshold of PIP_3_ necessary to achieve robust BTK activation in response to upstream B cell receptor (BCR) signaling events. Our liposome-based kinase assay showed a 2.77-fold increase in the extent of BTK autophosphorylation. Further mathematical modeling of the data corroborates with this change. Conceivably, at higher PIP_3_ levels where BTK can be fully saturated on the high-affinity PIP_3_, this effect of PS binding is expected to plateau out. This is also supported in our kinase assay, where no effect of PS concentration was observed at 5% PIP3 containing liposomes. At higher PIP_3_ concentrations, the excess of PIP_3_, making up to one in 20 lipids, begins to occupy all BTK, leaving less free BTK to recognize PS to recruit it on the surface, explaining how the difference in PIP_3_ concentration alters PS sensitivity.

An unresolved aspect of this study is the precise delineation of the binding sites for PS on BTK. Despite extensive mutagenesis efforts, we were unable to localize specific residues whose mutation disrupted PS binding. This finding raises the possibility that PS binding might not involve distinct, well-defined sites but rather general electrostatic surface interactions. Surface charge analysis further supports this hypothesis, demonstrating a large electropositive surface surrounding the canonical PIP_3_ site (Fig. 5A). Such an extensive electropositive surface is likely to facilitate a generalized electrostatic mechanism, enabling PS-associated membrane bound BTK to remain conformationally flexible, while gliding along the plasma membrane to present its canonical binding site optimally for high-affinity PIP_3_ interactions. This also guides the PHTH domain to have the electrostatically positive surface surrounding the canonical PIP_3_ site facing towards the membrane surface (Fig. 5B). This orientation can facilitate faster search of the negatively charged inner leaflet and have the PHTH domain already orientated with the area around the canonical site towards the lipid head groups (Fig. 5B). An electrostatic gliding mechanism might allow BTK to rapidly scan the PS-rich inner leaflet of the plasma membrane for PIP_3_. Indeed, previous computational and FRET-based studies on isolated PH domains of GRP1, a guanine nucleotide exchange factor, has proposed electrostatic hopping that can facilitate PIP_3_ binding (33). Notably, the nMS-based workflow provides the analytical precision required to unravel such nuanced multi-modal interactions directly from lipid membranes, as exemplified by our successful demonstration of dual PIP_3_ and PS binding to BTK. The autophosphorylation assay further illustrates how these weak-affinity electrostatic interactions significantly reduce the threshold concentration of PIP_3_ required to achieve physiologically relevant kinase activation upon BCR engagement. Given that the PH domain is one of the most widespread membrane-binding domains in the human proteome, found in over 250 distinct proteins, understanding how cells rapidly recruit PH-domains proteins using short-lived low abundant lipid such as PIP_3_ presents an exciting mechanistic question (53). Our work establishes a broadly applicable and robust analytical framework for investigating multi-modal peripheral membrane protein-lipid interactions. The nMS approach described herein, using a highly clinically-relevant peripheral membrane protein, offers unprecedented molecular resolution, enabling the identification of novel protein-lipid interactions, and provides a powerful tool for understanding the fundamental principles governing the selective recruitment of cytosolic effectors to various cellular membranes.

**Figure 5.**
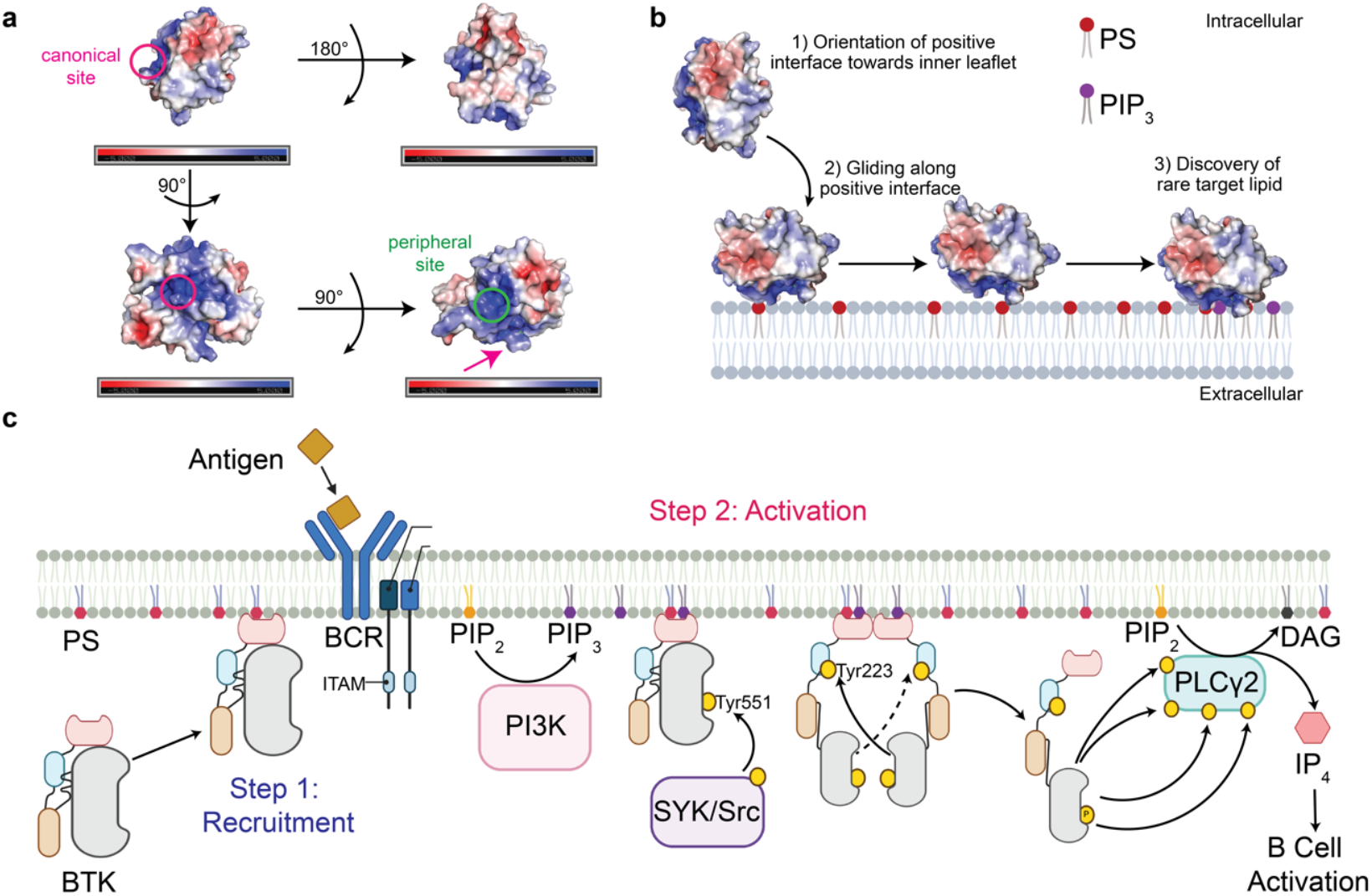
**(a) (top left)** Electrostatic map of the PHTH domain of BTK (PDB ID: 4Y94) (18). Electrostatics calculated on a single PHTH monomer using Pymol APBS electrostatics. Canonical site highlighted in fuschia circle **(top right)** Same plot orientated 180° around the x axis showing lack of electropositive character on this face. **(bottom left)** Plot from **top left** oriented 90° around the y axis, with the canonical PIP_3_ binding pocket highlighted in a fuschia circle. Large electrostatic positive character and pockets surround the canonical site. **(bottom right)** Plot from **bottom left** oriented 90° around the x axis highlighting another large electrostatic positive surface centered around the peripheral PIP_3_ binding site. Canonical site highlighted at the edge of the structure with a fuschia arrow. Though the sites are two hotspots for binding PIP_3_, many electropositive lysines and arginines are exposed on proximal loops to these sites as seen in **bottom left** and **bottom right** though have not been directly implicated in coordinating PIP_3_ through the two defined sites. **(b)** Electrostatic map shown in **(a)** and model of membrane search mechanism, showing orientation of electropositive character surrounding the PIP_3_ binding site gliding along membrane, orienting PHTH domain towards the anionic lipids. **(c)** Schematic depicting the proposed two-step binding of BTK on the inner leaflet of the plasma membrane. First, due to a low-affinity interaction with the highly abundant PS, BTK is recruited to the surface of the plasma membrane inner leaflet to dynamically explore for a high affinity activating ligand. Upon BCR stimulation, this initiates a cascade that activates two critical upstream proteins: PI3K and Src-family kinases. PI3K converts lipid phosphatidyl inositol (4,5) phosphate (PIP_2_) into phosphatidyl inositol (3,4,5) phosphate (PIP_3_). BTK is then activated by this high-affinity lipid and trans-autophosphorylates at Tyrosine 223 in the SH3 domain. Due to priming of BTK in the proximity of the membrane due to its recognition of PS, this activation can take place on a much faster scale. Src-family kinases, including LYN and FYN can phosphorylate Tyrosine 551 in the activation loop of BTK, increasing its catalytic activity. This fully active BTK can phosphorylate phospholipase C γ2, leading to cleavage of PIP_2_ to diacylglycerol and IP_3,_ a secondary messenger leading to downstream BCR activation.

## Materials and Methods

### Purifications

Plasmids were transformed into *Escherichia Coli* BL21 (DE3) cells for protein expression. For full-length BTK, *E. Coli* cells also containing the expression constructs for tyrosine phosphatase (YopH) and GroEL/ES were generated and used. *Bos Taurus* BTK was used, as it retains > 98% sequence homology and retains identical lipid binding sites and can be expressed in high quantities in *E. Coli*. Cells were grown in Terrific Broth (TB) with 70 µg/ml Kanamycin (with an additional 50 ug/ml Spectinomycin and 50 ug/ml Chloramphenicol for full-length BTK) to an optical density of 1.6 at 37° C, at which point an equivalent amount of 4° C TB with 70 µg/ml Kanamycin (with an additional 50 ug/ml Spectinomycin and 50 ug/ml Chloramphenicol for full-length TB), 100 µM ZnCl2, and 1 mM IPTG was added to induce a 16-hour expression at 18 C. Cells were harvested at 4,000 x g for 30 min and cell pellets resuspended in Ni-NTA buffer A (25 mM Tris pH 8.5, 500 mM NaCl, 5% glycerol and 20 mM imidazole) with protease inhibitor cocktail and DNAse. Cells were lysed via sonication (40% amplitude, 20 sec on, 45 sec off, 5 min total) and debris centrifuged out at 200,000 x g for 45 minutes at 4° C. The supernatant was loaded onto an HisTrap 5 mL column using a peristaltic pump at 1 mL/min and washed with 10 Column volume (CV) Ni-NTA buffer A. The protein was eluted via a gradient going from 0% Ni NTA Elution buffer (25 mM Tris pH 8.5, 500 mM NaCl, 5% glycerol and 250 mM imidazole) to 100% over 5 CV increasing by 20% B for each CV. The eluted fractions containing protein were equilibrated into Desalting Buffer (25 mM Tris pH 8.0, 400 mM NaCl) on a HiPrep 26/10 desalting column to remove imidazole. The protein was incubated overnight with 500 µg His-tagged ULP1 protease at 4° C to cleave the His-SUMO tag leaving the untagged protein and the cleaved His-SUMO and His-ULP1. Cleaved protein was then flowed over a HisTrap 5 mL column equilibrated in Ni-NTA buffer A to remove ULP1 and the His-SUMO product and collected in the flow-through. The flowthrough containing untagged protein was purified further using size exclusion chromatography (SEC) into SEC buffer (25 mM Tris-HCl, pH 8.0, 400 mM NaCl, 5% glycerol, and 1.5 mM TCEP) on a HiLoad 16/600 Superdex 200 pg column prior to concentration, aliquoting and flash-freezing.

### Preparation of samples for nativeMS

For native mass spectrometry of the purified proteins, proteins were buffer exchanged into 500 mM ammonium acetate, 1 mM DTT using Zeba desalting columns (Thermo Fisher Scientific) at room temperature. Protein concentration was determined after exchange into buffer. Protein ranges were kept between 1 and 5 µM and electrospray performed using in-house nano-emitter capillaries. These were generated by pulling borosilicate glass (O.D. – 1.2 mm, I.D. – 0.69 mm, 10 cm, Sutter Instruments) using a Flaming/Brown micropipette puller (Model P-100, Sutter Instruments) to a tip diameter within 4-5 µm. Platinum wire was inserted into the emitter and connected to the source for sample charging. nMS was performed In a Q Exactive UHMR using a Nanospray Flex ion source (Thermo Fisher Scientific). Mass spectrometry parameters were optimized for each sample, for instance, spray voltage was in the range between 0.9 – 1.5 kV, the capillary temperature was 150-250 °C, the resolving power of the MS was in the range between 3,125 – 6,250 the in-source desolvation was between 50V and 300V. For incubation with lipids in detergent, lipids from Avanti polar lipids were dried under nitrogen flow and vacuum prior to resuspension via vortex in 500 mM ammonium acetate + 1 mM DTT + 1.25x CMC OGNG. Protein and lipid were incubated for one hour prior to being subjected to nMS analysis as described above. PHTH domain interactions with lipid were kept at 4 µM protein to 200 µM lipid. For the titration, FL BTK is kept at 2.5 μM, PS at 125 μM, and IP_4_ concentration at 0 μM, 0.625 μM, 1.25 μM, and 2.5 μM.

### Fluorescence Polarization

Samples were prepared in four replicates for each condition using phosphatidylserine 18:1-12:0 labeled on the fatty-acyl chain with nitrobenzodiazole (NBD) from Avanti Polar Lipids. NBD-PS was dried under Nitrogen gas and vacuum to remove residual solvent before being resuspended in Polarization buffer (25 mM Tris pH 8.0, 400 mM NaCl, 1.5 mM TCEP, 5% glycerol and 0.029% OGNG). NBD-PS at 25 nM was used in all conditions. OGNG kept at half the critical micellar concentration to provide solubilization without forming larger micelles that could interfere with solution dynamics. Increasing concentrations of the purified bovine wild-type PHTH domain were added at intervals between 0 and 200 µM, incubated for 30 minutes and binding measured by fluorescence polarization in black-welled 96-well plates. Final buffer conditions in all wells were Polarization buffer and samples were measured using an Agilent BioTek Synergy H1. Linearly polarized light at 485 nm was sent through to excite the NBD and the parallel and perpendicular emitted light at 528 nm was measured, read at a 6.25 mm height. Filter settings were Excitation 485/20 and Emission 528/20. Polarization values were calculated using the instrument software Gen5 v3.11 following the standard formula P = (F|| - F⊥)/(F|| + F⊥). Briefly, this is the difference between the emitted intensity parallel to the excitation light plane and perpendicular over the total emitted intensity. Polarization values are reported in mP (milliPolarization units).

### Liposome Formation

Lipids of appropriate concentrations were dried under nitrogen and vacuum for one hour before being resuspended in 500 mM ammonium acetate supplemented with 1 mM DTT on vortex for one hour. Texas Red DHPE as a minor component was included to monitor vesicle stability during the extrusion process for nMS experiments. Lipid mixtures were then subjected to ten freeze-thaw cycles and extruded using an Avestin liposofast extruder with 100 nm membranes to ensure homogeneity. BTK was prepared in 500 mM ammonium acetate and 1 mM DTT as described previously. BTK and respective liposomes were incubated at concentrations of 1.775 µM and 1.5 mM liposomes in 500 mM ammonium acetate and 1 mM DTT. For the detection of BTK in the presence of both liposomes and IP4, 0.85 µM IP_4_ was supplemented. IP_4_ was diluted to a working concentration in the same buffer prior to addition.

### Kinase Activity

Liposomes were dried under nitrogen and vacuumed, resuspended in liposome resuspension buffer (25 mM Tris-HCl pH 7.4, 100 mM NaCl) at 2 mM and subjected to ten free-thaw cycles using a 50° C water bath and liquid nitrogen. These hydrated vesicles are then extruded to ensure 100 nm homogeneity using an Avestin LiposoFast. Full-length BTK at 4 µM and liposomes at 500 µM were prepared in kinase buffer (25 mM Tris pH 7.5, 150 mM NaCl, 5% glycerol) and subjected to a ten-minute incubation at room temperature. At which, time point 0 was taken. At time point 0, an equal volume of kinase reaction buffer (25 mM Tris pH 7.5, 150 mM NaCl, 5% glycerol, 20 mM MgCl2, 2 mM ATP, and 2mM sodium orthovanadate) or kinase negative control buffer (25 mM Tris pH 7.5, 150 mM NaCl, 5% glycerol and 2mM sodium orthovanadate) was added and the reaction proceeded for five minutes. All kinase and negative control reactions were stopped via the addition of an equivalent volume of 2x Laemmli sample buffer (Bio-Rad) with 100 mM EGTA and 5% β-mercaptoethanol and 15 minutes of 95 C boiling.

*Auto-*phosphorylation of BTK at phosphoTyr223 was measured using western blotting. Briefly, samples were run on an SDS-PAGE gel, before transfer to a PVDF membrane using a Tank transfer apparatus (BioRad). Membranes were blocked for one hour rocking at room temperature in 4% Bovine Serum Albumin in Tris-buffered saline with tween (TBST). Overnight incubation in a 4 C rocker with 1:2000 (1:500 for 5% PIP_3_) phosphoTyr223 antibody in 1% BSA in TBST was followed by five ten-minute TBST washes and secondary incubation with 1:5000 (1:10,000) goat anti-mouse in 1% BSA in TBST and a sequential five ten-minute TBST washes prior to imaging with SuperSignal West Femto or Pico PLUS Maximum Sensitivity Substrate (Thermo Fisher Scientific). For Coomassie staining, the gels were placed into fixation solution (40% methanol, 10% acetic acid), microwaved for thirty seconds and placed on a room temperature rocker for thirty minutes. Then, the gels were microwaved for thirty seconds in QC colloidal Coomassie stain (BioRad) and placed on a room temperature rocker overnight before rinsing several times with water for destaining. Both chemiluminescence and Coomassie blue staining were imaged on a BioRad Chemidoc. Pre-stained ladder was run for molecular weight conformation and imaged in the westerns. Analysis and integration of the bands was performed on Fiji.

### Mathematical modeling

Mathematical model adopted from Qi et. al. (2015) (18). Previous estimates of effective BTK concentration (C) on the membrane surface were calculated as follows:

Firstly, the active volume where BTK can operate within the lipid vesicle context, or the shell of the vesicle containing BTK is calculated as V = A x L (SI Fig. 8A). In the model, A is the surface area of the vesicle and L is the length of BTK, around 100 Å as in the BTK composite full-length structure. From this, C can be calculated as 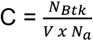 where N_BTK_ is the number of BTK on a vesicle and N_a_ is Avogadro’s number. N_BTK_ is calculated from the molar fraction (*f*) of PIP_3_ and the total number of lipids in the vesicle (N_0_). N_0_ is given by A x A_0_, the surface area of a lipid head group, at roughly 0.6 nm^2^(54).

C can then be calculated as:

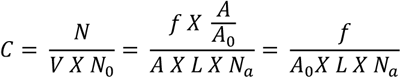

From here, we can modify this equation for our kinase assays, to become

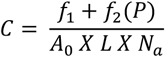

Where f_1_ represents molar fraction of PIP_3_, f_2_ represents molar fraction of PS and P represents the percentage of PS occupied by BTK. We then estimate P (the fraction of PS bound) based on our experimental data, and assuming the bulk of IP_4_ (PIP_3_ analog) is bound in the 1:1 stoichiometry as in the mathematical model. In our experimental titration with 0.625 µM IP_4_ we are using exactly 200-fold the concentration of PS as IP_4_ and achieving roughly 2-fold (1.916) the area of bound BTK. This approximates to around 1 in 100 PS is bound to BTK using the PIP_3_ analog for control. Our values become f_1_ = 0.005, 0.01, 0.025 and 0.05 according to PIP_3_ concentrations, f_2_ = 0, 0.1 and 0.2 accordingly P=0.01 for 1 in 100 bound, A_0_ = 60 Å^2^, N_a_ = 6.03 x 10^23^ mol^-1^.

## Acknowledgments

We would like to thank Dr. John Kuriyan and his lab members for sharing the BTK plasmids and Dr. Neel Shah for the YopH construct. We would like to thank Titus Boggon and his lab for the assistance with their plate reader for the fluorescence polarization assays. This work was supported by the National Institutes of Health grant R01GM141192 (to K. G.), R35GM147095 (to M.B.) and F31CA278383 (to R.M.).

