## Supplementary material for "Two-Step Mechanism of Bruton’s Tyrosine Kinase Membrane Recruitment and Activation": SI

**This PDF file includes:**

Figures S1 to S7

**Figures**

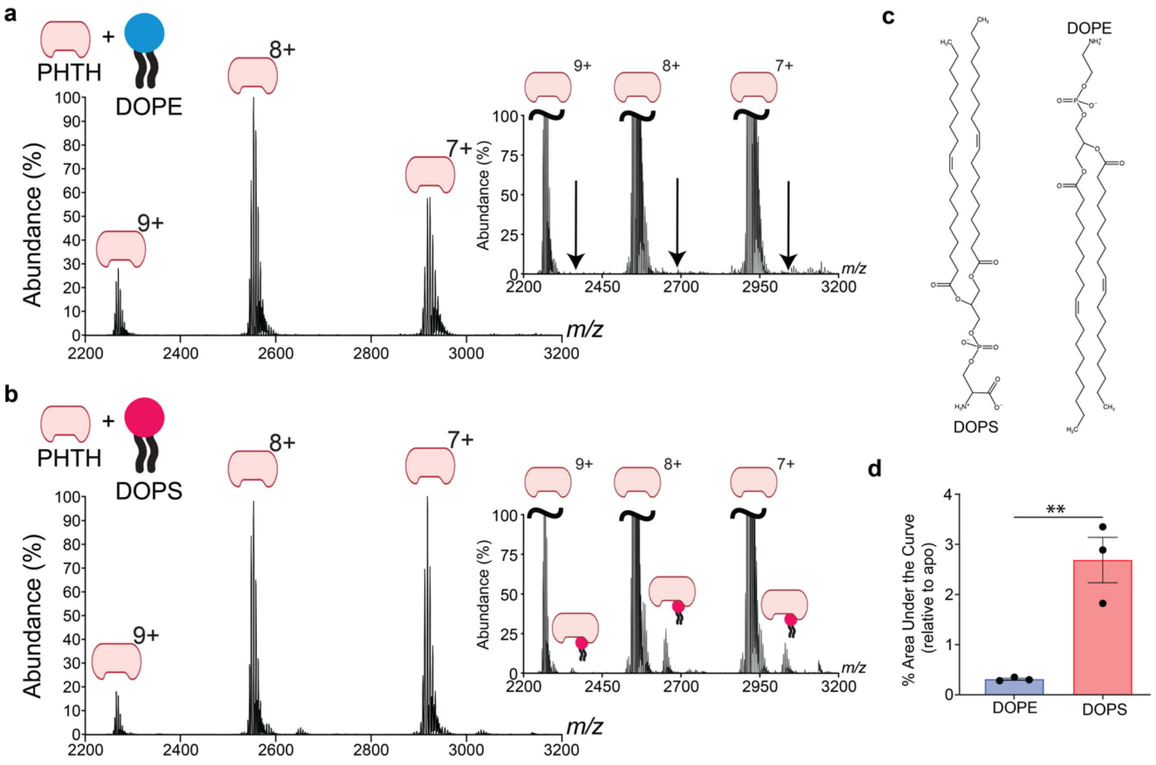


**Fig. S1. PHTH domain specificity of PS. (a)** Native mass spectra of the isolated PH-TH domain after incubation with 18:1-18:1 phosphatidylethanolamine (DOPE). No PE-bound peaks are identified. Arrows indicate predicted *m/z* for PE-bound peaks. Inset shows normalization to 10% of primary peak. **(b)** Native mass spectra of the isolated PHTH domain in the presence 18:1-18:1 phosphatidylserine (DOPS) under the same conditions reveals specific PS binding. Inset shows normalization to 10% of primary peak. **(c)** Schematics showing the structure of DOPS (left) and DOPE (right). The phospholipid backbones are identical, indicating any difference in behavior is specific to the head group. **(d)** Quantification of the lipid binding, shown as area under the curve relative to the apo 8+ charge state (mean ± sem, n=3). Statistics calculated using unpaired, nonparametric t-test on Graphpad Prism.

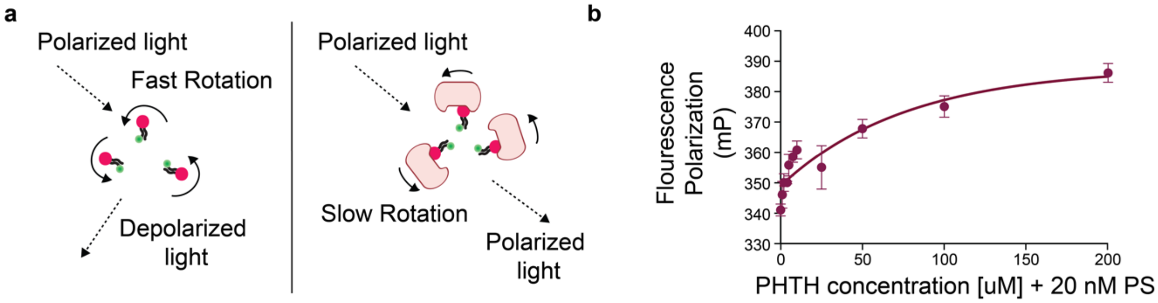


**Fig. S2. Fluorescence Polarization of PHTH and NBD-PS. (a)** Schematic depicting the mechanism of fluorescence polarization. Left: NBD-labeled PS in solution. Polarized light at wavelength 485 nm is used to excite the sample. Due to fast tumbling in solution of a small molecule, the emitted light at 528 nm is largely depolarized. Right: Binding of the much larger PHTH domain to the NBD-PS slows the rotation of the NBD-PS, reducing the depolarization of emitted light. **(b)** Chart showing the fluorescence polarization of NBD-PS as a function of PHTH domain concentration. Fluorescence millipolarization (mP) units measure the ratio of perpendicular/parallel light (see methods). Each concentration point was performed in replicate (mean ± sem, n=4). The increase in polarization as PHTH domain correlates with the binding and slower rotation of the PS.

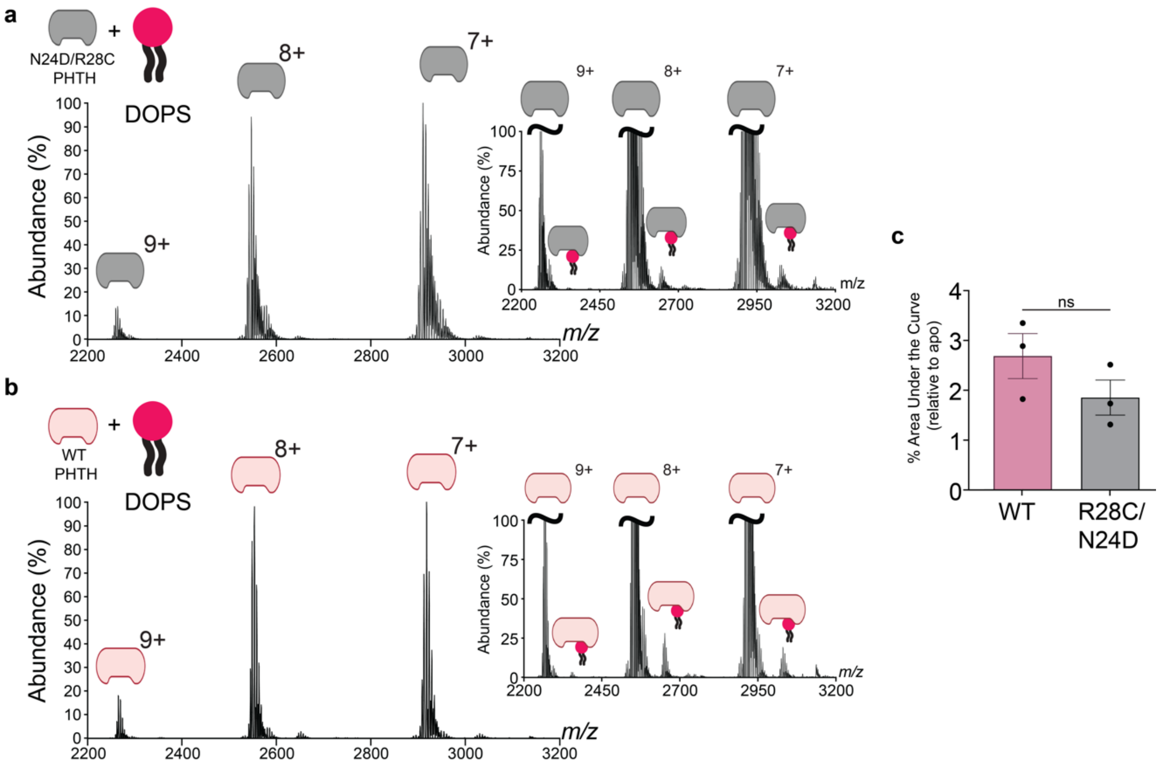


**Fig. S3. Canonical site (R28C/N24D) mutant PHTH domain retains PS binding: (a)** Native mass spectra of the isolated R28C/N24D mutant PHTH module in the presence of 18:1-18:1 phosphatidylserine (DOPS). Inset shows normalization to 10% of primary peak. **(b)** Native mass spectra of the isolated wild-type PHTH module in the presence of DOPS. Conditions are the same as in (a). Inset shows normalization to 10% of primary peak. **(c)** Area under the curve calculated relative to the primary charge state (8+) (mean ± sem, n=3). Statistics calculated using unpaired, nonparametric t-test on Graphpad Prism.

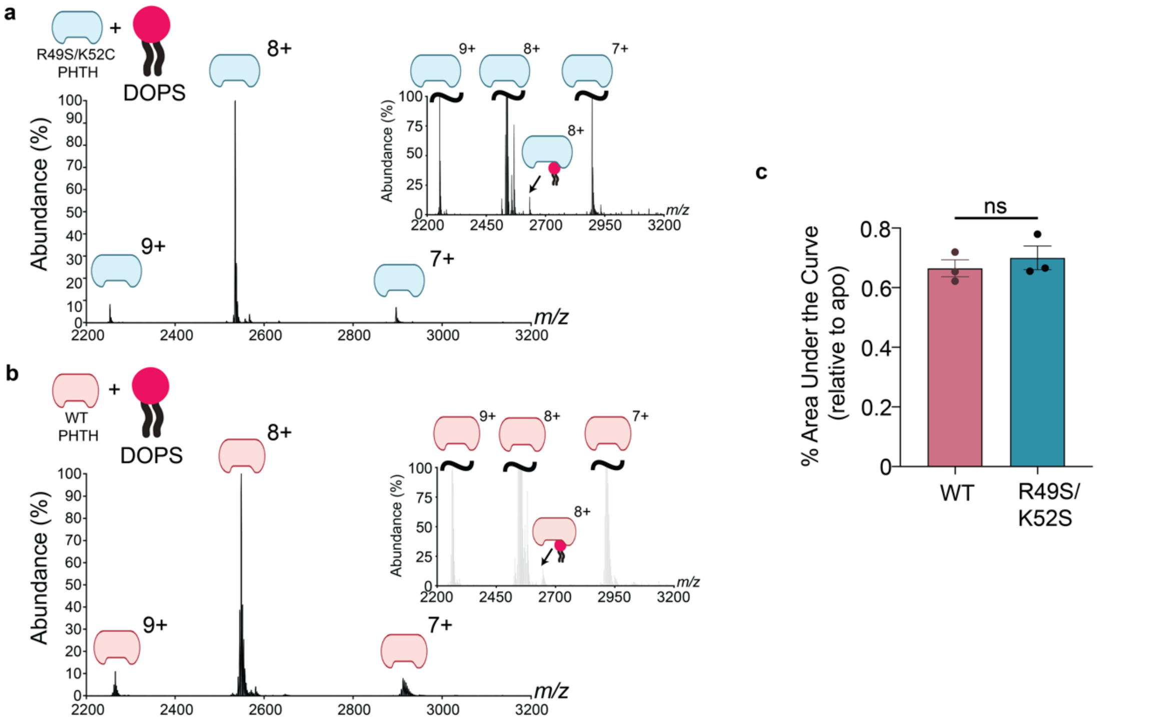


**Fig. S4. Peripheral site mutant PHTH domain retains PS binding**: **(a)** Native mass spectra of the isolated K49S/R52S mutant PHTH module in the presence of 18:1-18:1 phosphatidylserine (DOPS). Inset shows normalization to 5% of primary peak. **(b)** Native mass spectra of the isolated wild-type PHTH module in the presence of DOPS. Conditions are the same as in (a). Inset shows normalization to 5% of primary peak. **(c)** Area under the curve calculated relative to the primary charge state (8+). The area under the curve for each lipid-adducted species is calculated as a percentage of that same apo peak (mean ± sem, n=3). Statistics calculated using unpaired, nonparametric t-test on Graphpad Prism.

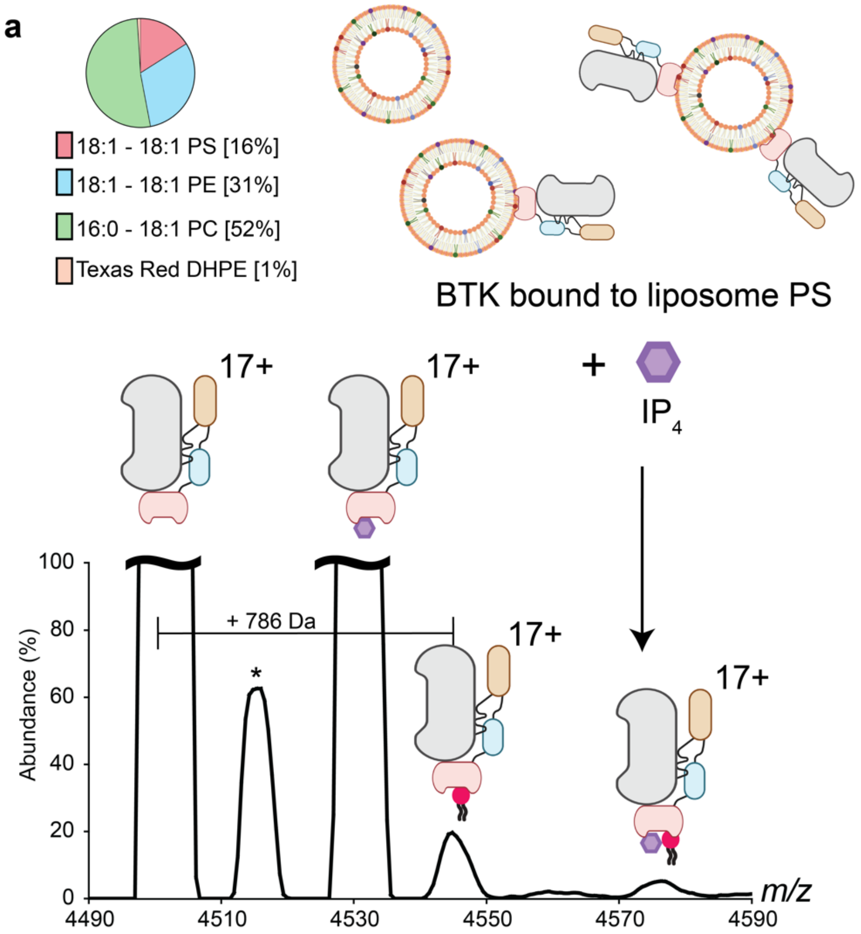


**Fig. S5. BTK bound to membrane PS and IP$_4$: (a)** nMS of Btk in the presence of liposomes containing PC/PE/PS and IP$_4$. Spectra highlights the binding of BTK to IP$_4$ and PS both individually and in unison.

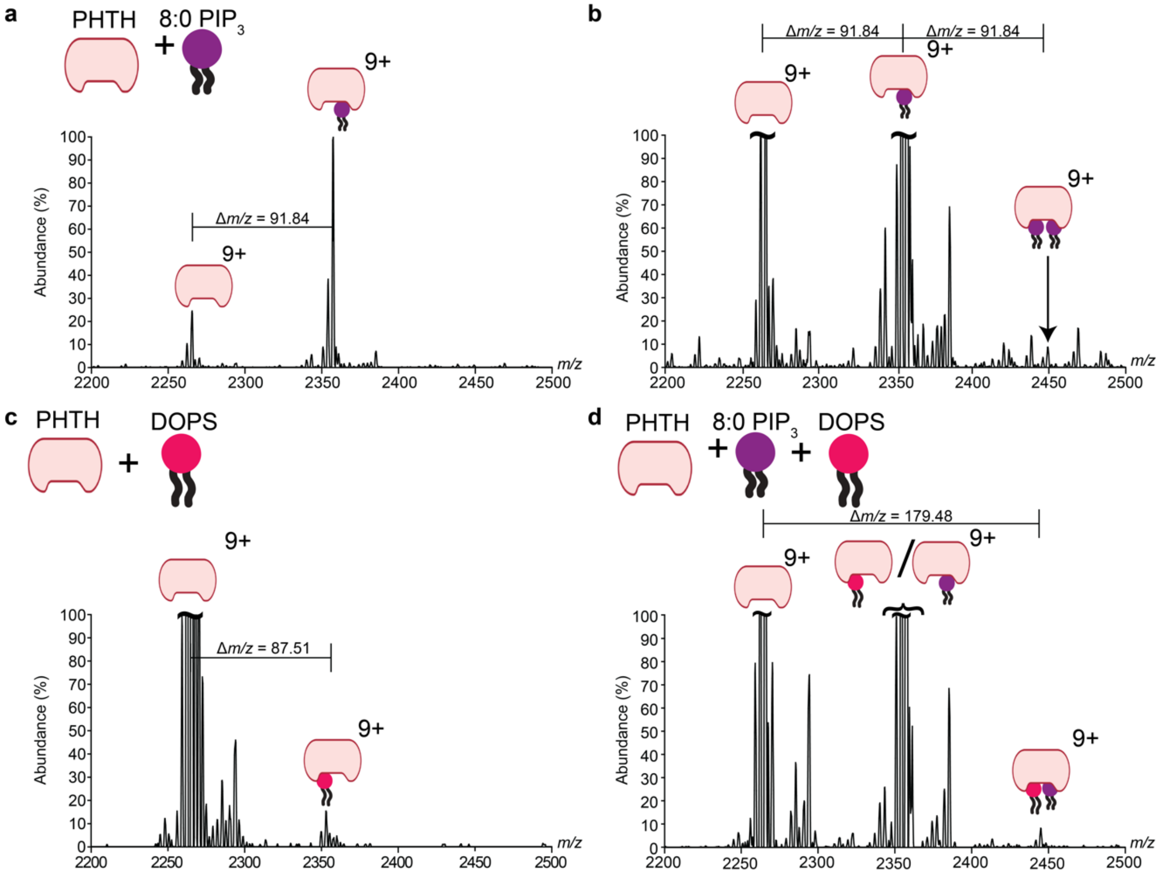


**Fig. S6. Supplemental to dual binding of BTK PHTH to PS and $PIP_3$. (a)** PHTH domain in the presence of 8:0 $PIP_3$ only in excess. The mass addition of 8:0 $PIP_3$ leads to a *m/z* difference of 91.84 (2357.43 – 2265.59) between the apo and lipid-bound charge states. **(b)** PHTH domain in the presence of 8:0 $PIP_3$ only normalized to 10% of highest peak in the spectra. Binding of a second 8:0 $PIP_3$ species reveals a peak at 2449.27 *m/z*, at another addition of 91.84 *m/z.* **(c)** PHTH domain in the presence of DOPS only normalized to 10% of the base peak in the spectra. The mass addition of DOPS leads to a *m/z* difference of 87.51 (2353.07 – 2265.56) between the apo and lipid-bound charge states. **(d)** PHTH domain in the presence of 8:0 $PIP_3$ and PS normalized to 10% of the base peak in the spectra. A peak exists at 2445.07 *m/z* at a 179.48 *m/z* addition to the apo peak. The peak corresponding to two 8:0 $PIP_3$ bound simultaneously would be at 2449.27 *m/z* as shown in b. The predicted *m/z* for two DOPS simultaneously bound would be at 2440.61 *m/z*. The actual peak (*2445.07 m/z*) aligns with one of each lipid being bound.

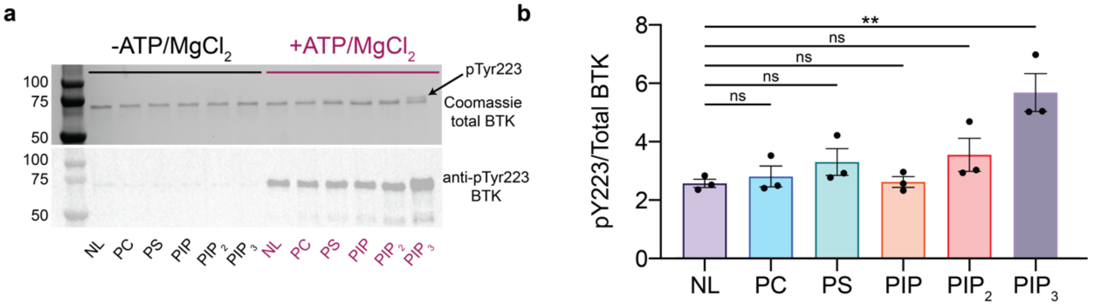


**Fig. S7. Anionic Lipids alone do not activate BTK. (a)** Kinase assay to demonstrate the effect of anionic lipids in the absence of $PIP_3$ on the kinase activity of BTK. Liposome lipid concentrations are NL – No liposome, PC – 100 mol %, PC/PS – 60/40 mol %, PC/PS/PIP – 87/10/3 mol %, PC/PS/PIP2 – 88/10/2 mol %, PC/PS/PIP3 – 88.5/10/1 mol %. **(top)** Representative Coomassie stain of each condition to demonstrate total Btk concentration in both the negative control in the absence of ATP and $MgCl_2$ as well as in the reaction conditions. The upper migrating band indicating the presence of phosphorylation in the 1% $PIP_3$ is highlighted with an arrow and annotated as pTyr223. **(bottom)** Representative anti-phosphoTyr223 BTK western. **(b)** Quantification of the BTK phosphoTyr223 relative to total BTK in each condition. All data shown as mean ± sem from n = 3 independent replicates. Results shown as unpaired parametric t-test. P-values plotted in a separate table. Statistical significance defined as p>0.05, ns; p< 0.05 as *, p<0.01 as ** and p<0.001 as ***.

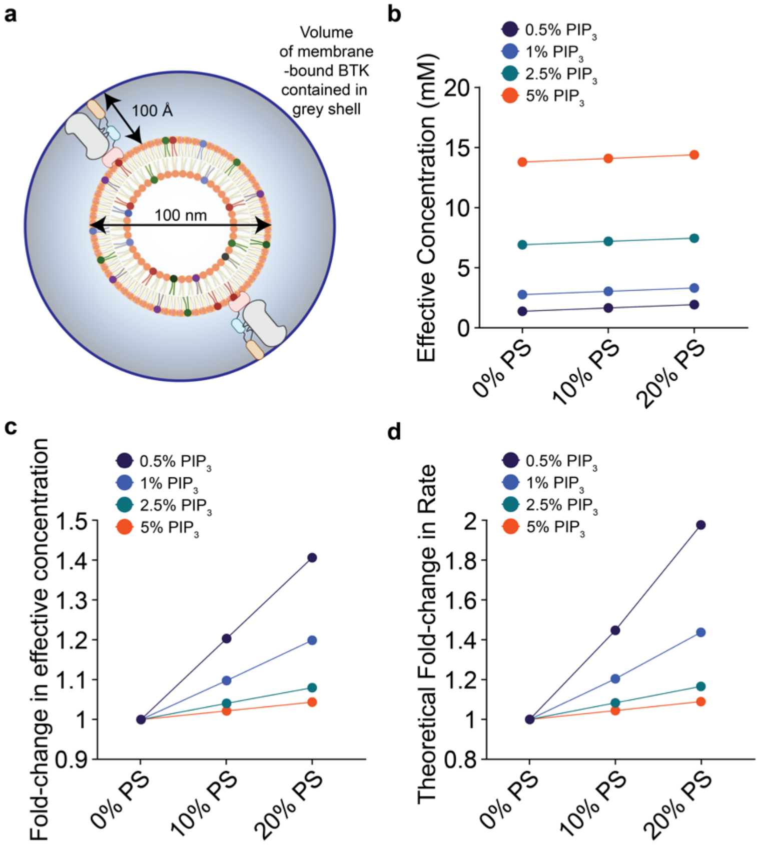


**Fig. S8. Mathematical model for membrane-bound BTK concentration. (a)** Model demonstrating how volume is calculated using the diameter of the lipid vesicle and the predicted length of full-length BTK. **(b)** Changes in the calculated effective concentration using the mathematical model for membrane-proximal BTK as a function of a set $PIP_3$ and changing PS concentration. As the PS concentration changes, the mM change is consistent. **(c)** Changes in the fold-change of effective concentration relative to 0% PS for each $PIP_3$ concentration. At lower $PIP_3$ concentrations, this leads to a more drastic change in the effective concentration. **(d)** Theoretical fold-change in rate calculated as the square of the fold-change in effective concentration, proportional to $[BTK]^2$. This provides an even more drastic increase as we consider the [BTK] as both the enzyme and substrate in a kinase autophosphorylation reaction.
